# Investigating sensorimotor beta burst dynamics as a robust biomarker for graded force modulation in humans

**DOI:** 10.64898/2026.05.07.723396

**Authors:** Md Shaheen Perwez, James J. Bonaiuto, Bhivraj Suthar, Vignesh Muralidharan

**Author notes:** shared senior authors (Bhivraj Suthar, Vignesh Muralidharan).

## Abstract

The most prominent signature associated with motor execution and motor imagery is the event-related desynchronisation and synchronisation (ERD/S) in the mu and beta bands (8-30 Hz). In the context of brain-computer interfaces (BCI), this ERD/S signature is helpful for binary decisions, such as left vs. right imagery, but it is not a robust biomarker for continuous prediction, such as precisely decoding different levels of force application. This is essential for developing better BCI applications with precise dynamic force outputs. Recent studies have revealed that sensorimotor beta bursts have a stronger relationship with motor control, even at a single-trial level, than mu and beta frequency band power. We, therefore, investigated whether the transient nature of beta bursts provide an alternative, but robust biomarker for BCI force decoding. Here, we designed an experiment where human participants (N = 16) performed both motor execution (ME) at four force levels (10%, 25%, 50%, and 75% of maximum voluntary contraction) and followed by imagination of the same, i.e. a motor imagery (MI) task, as their electroencephalogram (EEG) was recorded. We observed a clear and classical ERD pattern in the motor cortex region during the ME task, whereas it was less pronounced during the MI task. To explore the robust biomarkers for explanation of the force modulation we moved beyond classic trial average ERD/S and analysed the role of beta burst. After extracting sensorimotor beta bursts, we observed differences in spectral burst features between motor execution and imagery including burst amplitude, spectral width, and temporal width. Moreover, different force levels were correlated with changes in the burst amplitude and burst spectral width, specifically during motor execution. Interestingly, we found that different beta burst waveforms are associated with the different force levels and conditions. This suggests that the bursts-level features could be driven by changes in the underlying beta burst waveforms. Overall, our study shows that sensorimotor beta burst can be an important piece of the puzzle to implementing precise force control in brain-computer interface based prosthetics.

## Introduction

Decoding force modulation using electroencephalography (EEG) is an emerging area of brain-computer interface (BCI) research that aims to expand the command capabilities of BCI systems beyond simple binary control, enabling more sophisticated interaction with external devices. This development is particularly relevant for achieving nuanced control over neuroprostheses, rehabilitation robots, or exoskeletons (Ding et al., 2025; Meng et al., 2016; Tang et al., 2016; Volkova et al., 2019; Wang et al., 2017). Force modulation in BCIs involves decoding kinetic parameters, such as grasp force, from neural signals to enable precise control of devices such as prosthetic hands (Ding et al., 2025; Edelman et al., 2016; Meng et al., 2016; Tang et al., 2016; Volkova et al., 2019). While invasive intracortical recordings (ECoG) have revealed that force-related signals exhibit a dynamic, nonlinear relationship with motor cortex activity (Acharya et al., 2010; Nakanishi et al., 2014; Okorokova et al., 2024), translating these findings to non-invasive modalities such as EEG-based systems remains an active area of research (Chakarov et al., 2009; Fry et al., 2016; Haddix et al., 2021; J. Z. Liu et al., 2005; Nakayashiki et al., 2014; Slobounov et al., 2002).

Studies show that imagining or attempting force exertion modulates EEG patterns, particularly in sensorimotor areas, with stronger event-related desynchronization (ERD) at higher force levels in mu and beta rhythms (Alayrangues et al., 2019; Pfurtscheller, 1981; Pfurtscheller et al., 1996; Pfurtscheller & Aranibar, 1977; Pfurtscheller & Lopes da Silva, 1999; Wang et al., 2017). While machine learning can successfully decode basic binary motor imagery in stroke patients (H. Liu et al., 2024), expanding these models to decode continuous force remains challenging due to pathological neural signatures and the reliance on simple binary command setups (Barone & Rossiter, 2021; Haddix et al., 2021; Peter et al., 2022). Single-trial EEG recorded during motor imagery of two isometric grip force levels (5% vs. 40% MVC) can be reliably classified using combined common spatial pattern and coherence features with a Support Vector Machine (SVM), achieving a mean offline accuracy of 81.4% and enabling real-time control (Hualiang et al., 2023) . Furthermore, in the same study, participants successfully used this force-level imagination paradigm to play an online ball-movement game, achieving a mean accuracy of 76.7%, validating its potential to provide continuous, graded commands to noninvasive BCIs (Hualiang et al., 2023). Force modulation extends motor imagery (MI) paradigms, in which users imagine varying force loads (e.g., 10% vs. 30% of maximum voluntary contraction), thereby enhancing BCI command sets for rehabilitation applications (Wang et al., 2017). Advanced algorithms, including recurrent neural networks, show promise for capturing temporal dynamics in force-related neural patterns (Okorokova et al., 2024), though their application to EEG-based force decoding requires further investigation.Currently, decoding continuous force application using EEG lies mostly in trial-averaged band-power features, such as mu and beta ERD (Fry et al., 2016; Wang et al., 2017). In addition to spectral power, signal complexity also tracks force output; for example, the fractal dimension (FD) of EEG signals over the motor cortex increases linearly by 10–20% as handgrip force rises from 20% to 80% MVC (J. Z. Liu et al., 2005). However, current decoding algorithms do not fully capture the nonlinear or transient dynamics underlying force modulation, thereby constraining real-time performance and adaptability.

Recent advances in our understanding of the neurophysiology of sensorimotor beta oscillations (13–30 Hz) show that they occur as transient, short-lived bursts rather than sustained oscillations (Bräcklein et al., 2022; Feingold et al., 2015; Little et al., 2019; Papadopoulos, Darmet, et al., 2024; Papadopoulos, Szul, et al., 2024; Rayson et al., 2022; Szul et al., 2023). These bursts have a wavelet-like shape in the time domain (Rayson et al., 2022; Szul et al., 2023) and are more spatially focal than temporally averaged beta amplitude (Little et al., 2019). In addition, their timing and amplitude reflect motor preparation and response dynamics (Little et al., 2019). In other studies, high amplitude and temporally closer sensorimotor beta bursts are directly linked to a significant increase in subsequent movement slowing ( Muralidharan & Aron, 2021). More recently, it has been shown that beta bursts do not have a stereotypical waveform, and their waveform shape can change based on context (Rayson et al., 2022; Szul et al.). This diversity suggests the possibility of modulating cortical sensorimotor beta bursts. However, given their transient nature, this modulation does not directly lead to fine-grained control over isometric force manipulation; instead, burst modulation acts inside a motor null-space relative to direct force production(Bräcklein et al., 2022; Churchland & Shenoy, 2024). Beta bursts parameters, specifically their amplitude and duration, systematically change across motor states such as rest, preparation, execution, and imagery, making them potentially superior to traditional spectral features for state classification. Furthermore, the execution of movement is specifically characterised by a significant reduction in both the amplitude and duration of these beta bursts compared to periods of rest or movement preparation (West et al., 2023). Contrary to the traditional view of continuous oscillations, both motor cortical beta activity and cortico-muscular connectivity actually manifest as short, transient bursts even when maintaining a sustained motor behavior (Echeverria-Altuna et al., 2022)

In this study, we recorded EEG data while participants applied different force levels, followed by imagining application of the same force levels. EEG data were then analysed for the task and force levels. Time–frequency analyses using superlet transform were performed to characterise oscillatory dynamics associated with motor execution and imagery, in the sensorimotor mu (8–13 Hz) and beta (13–30 Hz) bands. The superlet transform is an adaptive time-frequency decomposition technique based on Morlet wavelets to optimally balance temporal and spectral resolution (Moca et al., 2021). It achieves this optimal balance by linearly varying its order (a multiplier of the wavelet’s cycles) across the frequency range, making it highly sensitive for detecting transient neural bursts compared to traditional methods like the Continuous Wavelet Transform (CWT) which suffers from frequency smearing. In addition, beta-burst features were extracted to investigate transient oscillatory events underlying motor tasks with different force levels. The results revealed clear modulation of sensorimotor rhythms during motor execution, with stronger event-related desynchronisation and more pronounced beta-burst dynamics compared to motor imagery. In contrast, motor imagery elicited weaker patterns of oscillatory modulation, suggesting that although both conditions recruit overlapping neural mechanisms, the strength and temporal structure of cortical activity differ between executed and imagined force production. These findings provide insight into the neural representation of graded force which can be leveraged for EEG-based decoding of force levels, for instance, in brain–computer interface applications.

## Methods

### Participants and Setup

We had already established the number of participants (N=16) to collect EEG data in our preregistered hypothesis using G*Power (Faul et al., 2007) to calculate the required sample size (https://osf.io/rjbpt/files/a6nmw). We collected EEG data from 20 participants (10 females and 10 males; mean age = 22.2 years, range = 18–31 years). In 16 participants we observed a clear event-related desynchronization in the mu band (8–12 Hz) at C3 channel. These 16 participants were included for the analysis. All participants provided written informed consent in accordance with the Declaration of Helsinki. All procedures and protocols were approved by the Institutional Ethics Committee of the Indian Institute of Technology Jodhpur. All participants were right-handed and reported no known prior neurological disorders.

Participants applied and imagined different force levels while their hands were positioned in a custom-built isometric wrist-flexion dynamometer, which was securely fastened to the table in front of them **(Figure 1A)**. Initially, we measured each participant’s maximum voluntary force contraction by making them perform wrist flexion (Flexor carpi radialis muscle) **(Figure 1B)**. Following this, participants performed a practice session of the experimental paradigm (consisting of only two blocks) to become familiar with the task and practice force execution and imagery. No participants had prior experience with motor imagery.

**Figure 1.**
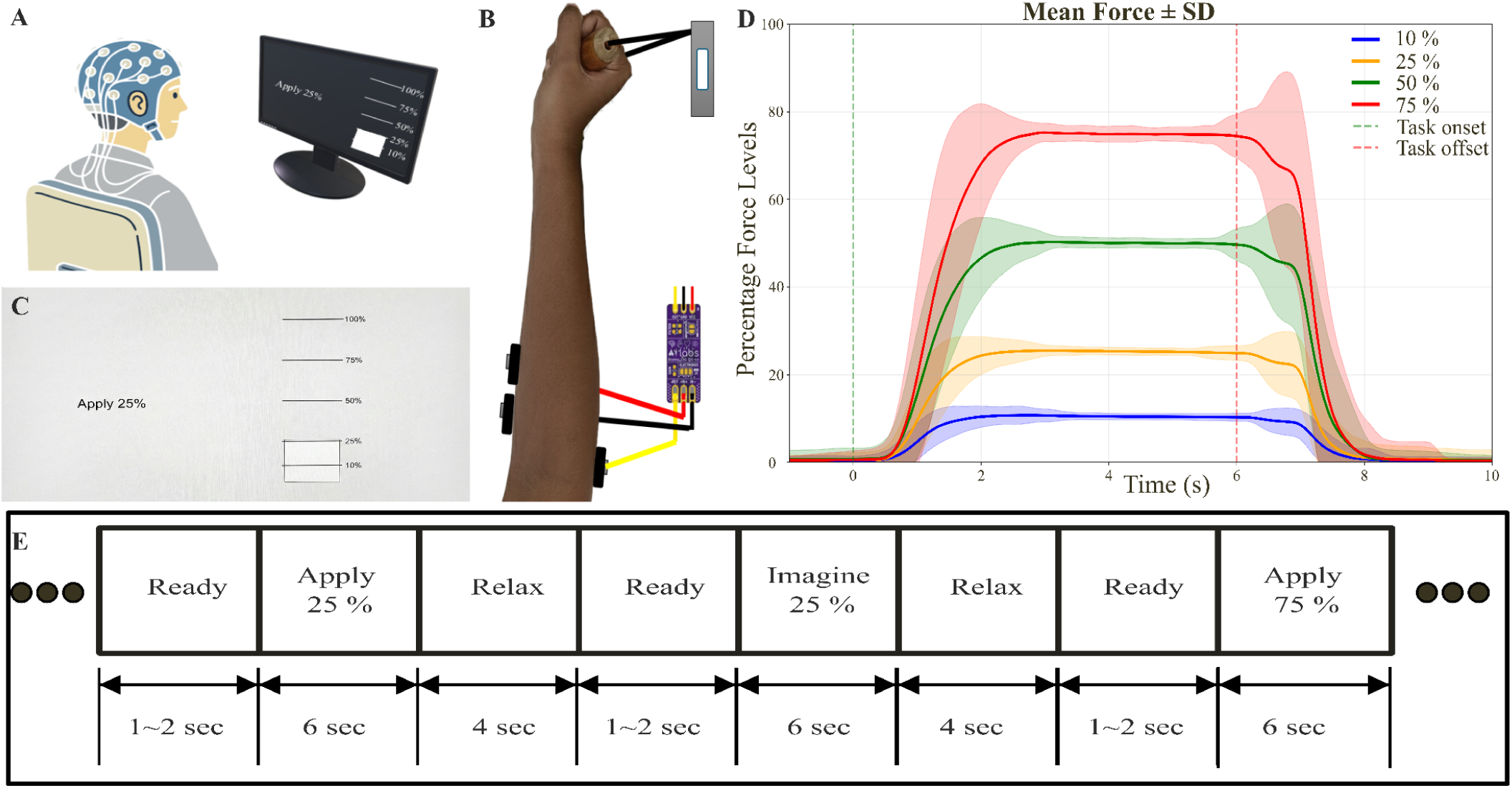
Experimental setup and force-control paradigm. **(A)** EEG recording setup: A participant seated in front of a computer display while wearing a 64-channel EEG cap (representative only) . Neural signals were recorded continuously during task performance as participants interacted with the force device using their dominant (right) hand. **(B)** Force measurement and EMG setup: Participants grasped a cylindrical handle instrumented with a force sensor, allowing for continuous measurement of the applied wrist flexion force. A bipolar surface EMG sensor was positioned over the forearm muscles (Flexor Carpi Radialis) to simultaneously record muscle activity during force production. Both the sensors’ outputs were sent to Arduino and temporally synchronised with the EEG acquisition. **(C)** Visual force-feedback interface presented to participants. Horizontal reference bars indicated the target force levels (10%, 25%, 50%, and 75% of each participant’s maximum voluntary force); participants were instructed to match and maintain the displayed target level (example shown for 25%). **(D)** Force traces illustrating the average force applied at different target levels (10%, 25%, 50%, and 75% of maximum voluntary force) during the motor execution task over time. Vertical green dashed line at t = 0 sec is the stimulus onset for different force levels, and the relaxation stimulus onset at t = 6 sec. Shaded region shows S.D. **(E)** Trial structure and timing of the experiment, showing the sequence of Ready, Apply/Imagine, and Relax phases and their respective durations.

### Experiment design

The experiment was conducted using PsychoPy Coder (Peirce, 2007). Participants were shown instructions on the left side of the screen, while a real-time bar plot of force levels was displayed on the right, marked at 10%, 25%, 50%, and 75% of the Maximum Voluntary Contraction (MVC) **(Figure 1C)**. During the experiment, the Motor Execution (ME) task was performed first, followed by the Motor Imagery (MI) task where the same force levels were imagined (**Figure 1E)**. Each trial commenced with a ready period lasting 1 to 2 seconds, followed by the application of one of the four force levels (10%, 25%, 50%, or 75% of MVC, **Figure 1D**) for 6 seconds, followed by a 4-second relaxation period. The bar plot continuously displayed the real-time force level applied by participants during the ME task. The MI task was similar to the ME task, but the participants had to imagine exerting the same force level. In this case, the feedback bar plot was not displayed. Each block consisted of 12 ME trials and 12 MI trials, with three trials per force levels per conditions, presented in a pseudorandomized order (**Figure 1E)**. Participants completed 10 blocks, totalling 30 trials per force level (ME and MI). After finishing each block, participants initiated the next block by pressing the spacebar, allowing them to rest as per their need before starting the next block.

### Data Acquisition (EEG, EMG and Force Sensor)

We recorded EEG data using a 64-channel BrainVision ActiCHamp amplifier and the BrainVision Recorder software with active electrodes (ActiCap snap cap, ActiCHamp Plus, Brain Products GmbH, Gilching, Germany). In this system, the electrodes are placed according to the international 10-20 system, with the reference electrode as FCz. We recorded the EEG at a sampling rate of 1000 Hz. Simultaneously, force was recorded using a 20 kg strain-gauge load cell arranged in a Wheatstone bridge configuration. The load cell output was digitised using a 16-bit ADS1115 analogue-to-digital converter interfaced with an Arduino microcontroller. The digitised signal was sent to the computer for real-time force-level plotting and simultaneous saving. The sensor was calibrated using known weights to convert raw ADC values into force unit . Force signals were recorded at a sampling rate of 250 Hz. EMG was acquired using the BioAmp ExG Pill (Upside Down Labs) at a sampling rate of 5 kHz. EEG, EMG and Force data were synchronised with the event markers for the task.

### Data Preprocessing

#### EEG

We preprocessed the recorded EEG data using the MATLAB-based EEGLAB toolbox (Delorme & Makeig, 2004) and custom scripts. The data were downsampled to 500 Hz and then filtered with a 1 Hz high-pass filter. Line noise was eliminated using the Zapline-Plus plugin (de Cheveigné, 2020). Subsequently, a 40 Hz low-pass filter and a common-average reference were applied. Further, we decomposed the EEG data using Independent Component Analysis (ICA) using the logistic Infomax ICA algorithm (Delorme & Makeig, 2004; Makeig et al., 1995). Artifactual components, including those associated with muscle activity, eye movements, and channel noise, were identified and removed manually . Then we used the DIPFIT plugin in EEGLAB to find the dipole that best represents the left sensorimotor cortex data. Then we extracted that IC data into a single-channel time series. For participants in whom we didn’t find a suitable IC representing the left sensorimotor cortex after dipole fitting, we used the C3 channel data (4/16 participants). Finally, the data were epoched from -1 s to 10 s relative to stimulus onset markers at the 10%, 25%, 50%, and 75% time points during the ME and MI tasks.

#### EMG

To preprocess EMG signals, a fourth-order Butterworth bandpass (75–150 Hz) zero-phase filter was applied, followed by notch filters at 100 Hz and 150 Hz to suppress residual line-noise harmonics. We then calculated the root-mean-square (RMS) of the filtered EMG using a rectangular window. Then, EMG was segmented into epochs time-locked to task-related event markers, spanning −1 s to +10 s relative to event onset. We then calculated the mean baseline activity from −1 to 0 s and the task-related activity from 2 to 6 s. The EMG ratio (active/baseline) was then calculated for each epoch to compare EMG activity during the ready period and the force application or imagery period, providing both trial-level and condition-level measures of muscle activation. In MI tasks we then excluded the trials with an EMG ratio (active/baseline) greater than 1.1. Across all participants on average 18.54 % of 10% MI tasks, 21.25 % of 25% MI tasks, 26.25 % of 50% MI tasks and 25.63 % of 75% MI tasks were rejected.

#### Force

Force analysis was performed on a trial basis which was time-locked to the task onset markers for different force levels (−1 to +10 s relative to event onset). For each participant, target force levels (10%, 25%, 50%, and 75%) were defined relative to the participant’s individual maximum voluntary force. Trial-wise force stability was assessed by comparing the force signal against condition-specific thresholds corresponding to the instructed force level. Only ME trials in which participants maintained the target force within ±10% of their participant-specific force level for at least 4 s were retained for further analysis, ensuring reliable and sustained force production. Across all participants on average 18 % of 10% ME tasks, 13.75 % of 25% ME tasks, 12.5 % of 50% ME tasks and 11.87 % of 75% ME tasks were rejected. The reach time and fall time of this stable force period were identified using threshold-crossing detection and were used to characterise force maintenance performance.

#### Data Analysis

As per our <u>OSF preregistration</u> ( https://osf.io/rjbpt/files/a6nmw ), we started by examining the left sensorimotor spatial filter derived from the ICA analysis (**Figure 2B)**. To extract beta bursts parameters (peak amplitude, peak frequency, peak time, full width at half maximum (FWHM) time, FWHM frequency, waveform) from the preprocessed epoched EEG data and for statistical analysis, we used MNE-Python (Gramfort et al., 2013).

**Figure 2.**
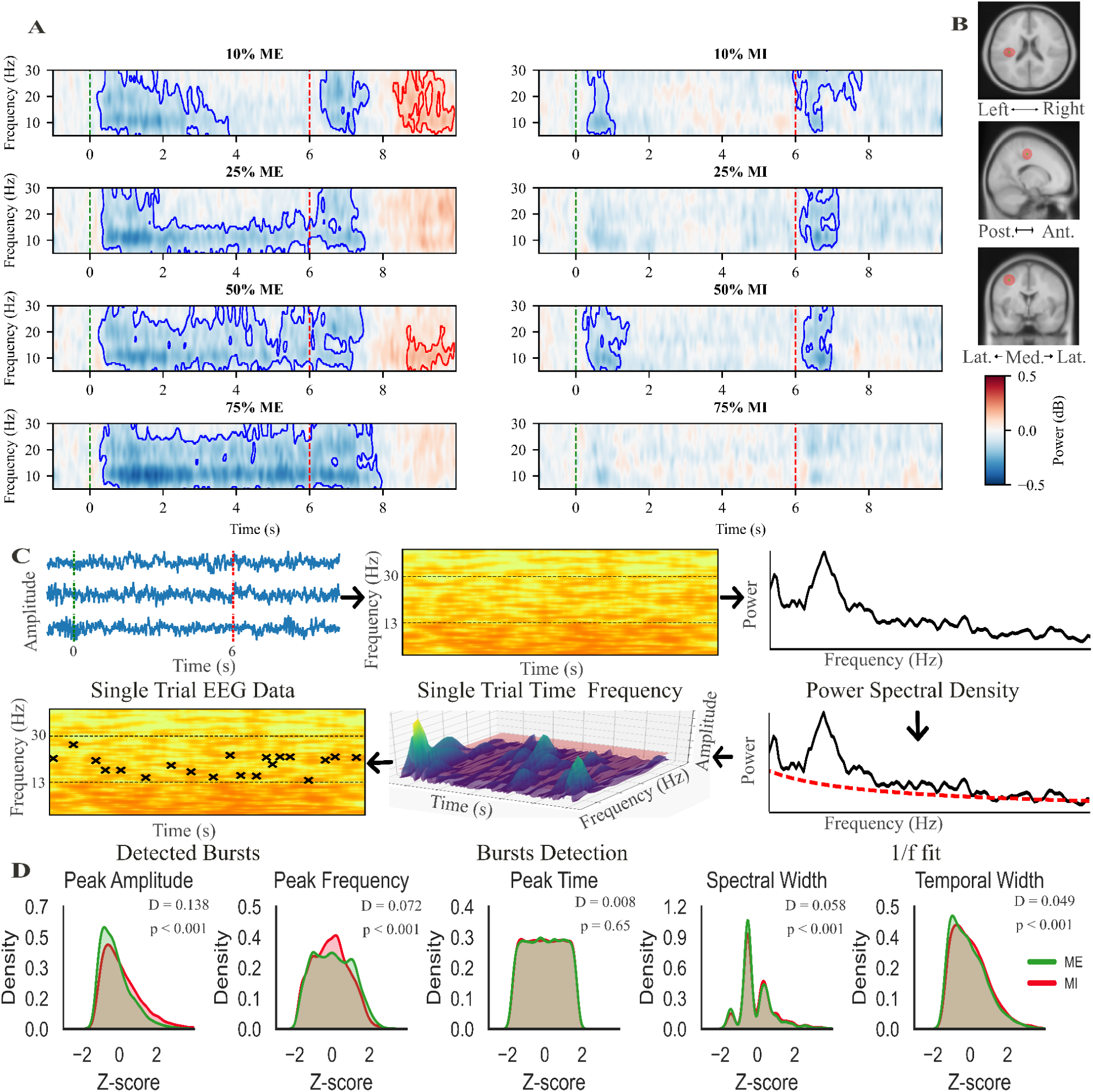
Mu–beta event-related desynchronization (ERD) during motor execution and motor imagery across force levels and extraction of bursts and their distribution in the ME and MI task. (A) ERDS: Time–frequency representations (5–30 Hz) show task-related changes in oscillatory power (log ratio, dB) relative to baseline (from t = -0.8 sec to t = 0 sec). Left panel: motor execution task; right panel: motor imagery task. Rows correspond to target force levels (10%, 25%, 50%, and 75% of individual maximum voluntary force). Negative values (blue) indicate ERD predominantly in the mu (8–13 Hz) and beta (13–30 Hz) bands during task engagement, with stronger and more sustained desynchronization during motor execution compared to motor imagery. The vertical green dashed line at t = 0 s marks task onset, and the vertical red dashed line at t = 6 s indicates task offset. Colour bars denote power change relative to baseline. Clusters overlaid with blue contours show ERD, and red contours show event-related desynchronization (ERS). **(B) Mean source localisation of left motor cortex activity:** Top view, sagittal view, and coronal view of the grand-average dipole location projected onto a standard anatomical brain, respectively. The green marker denotes the mean dipole position across participants, while the red shaded region represents the spatial dispersion (±1 standard deviation) of dipole locations, illustrating inter-participant variability. The dipole cluster is localised to the left motor cortex, consistent with sensorimotor involvement during the task. **(C) Beta Burst extraction:** We used the beta-burst extraction method as described by Szul et al. 2023. **(D) Bursts parameter distribution comparison between ME and MI tasks.**

#### Mu-beta ERD calculation for motor execution and imagination

Time–frequency power was estimated for each participant and condition using Morlet wavelets (1–40 Hz, 0.5 Hz steps; number of cycles = f/3). Since ERSP calculation is basically a macroscopic trial-averaged measure of sustained power, we didn’t implement the adaptive temporal resolution of the superlet transform which is best suitable to isolate highly transient bursts. Instead we used standard Morlet wavelets because their fixed-cycle approach provides the stable, uniform temporal smoothing needed to accurately capture broad, continuous amplitude trends across multiple trials without the need of too much computational power. Power was averaged across trials and baseline-corrected using a pre-stimulus interval of −0.8 sec to 0 sec with a log-ratio transform. Participant-level TFRs were then grand-averaged across participants. Statistical significance of task-related power changes was assessed using cluster-based one-sample permutation tests (Maris & Oostenveld, 2007). Time–frequency points exceeding a cluster-forming t-threshold corresponding to p = 0.05 were grouped into contiguous clusters across time and frequency. Cluster significance was then evaluated using 1000 permutations with random sign-flipping across subjects, and cluster-level p-values were obtained from the permutation distribution of the maximum cluster statistic, thereby controlling the family-wise error rate across the time–frequency space.

#### Time-resolved mu and beta band ERD - regression analysis

We applied a band-pass filter of 8-12 Hz for mu-band and 13-30 Hz for beta-band on preprocessed epoched EEG data from each participant and force condition, using a zero-phase 4th order Butterworth filter, and the analytic signal was obtained via the Hilbert transform to compute the instantaneous power envelope. Event-related desynchronization (ERD) was quantified on a trial-by-trial basis as percentage change relative to a stimulus onset baseline (−0.8 to 0 s). Trials were temporally realigned to the task reach time (time when the participants reached the instructed force levels) and segmented into a fixed window (−1 to 6 s). For each condition, ERD time courses were averaged across trials and smoothed using a Gaussian kernel (𝛔 = 20, i.e 40 ms). Group-level analysis was performed by pooling subject-wise ERD values across four force levels (10%, 25%, 50%, 75%). At each time point, a linear regression model was fitted to estimate the relationship between mu ERD and force level and beta ERD and Force Levels [**F=β_0_+β_1_**⋅**ERD(t)**], yielding a time-resolved regression coefficient (β₁) and associated p-value. Multiple comparisons across time were controlled using false discovery rate (FDR) correction (Benjamini–Hochberg, α = 0.05), and significant intervals were identified accordingly.

#### Beta-burst detection, burst parameter extraction, and Analysis

We extracted the beta bursts using adaptive-threshold method (Papadopoulos, Darmet, et al., 2024; Papadopoulos, Szul, et al., 2024; Szul et al., 2023). Single-trial time–frequency representations were computed using the adaptive superlet transform (1–40 Hz, 200 frequency bins). Beta burst detection was restricted to a search range of 10–33 Hz, with retained bursts required to have peak frequencies within 13–30 Hz. To account for the aperiodic (1/f) component of the spectrum, the average power spectral density across trials and time was parameterised using Specparam (Donoghue et al., 2020), and the resulting aperiodic fit was used to normalise the time–frequency data. Burst detection was then performed iteratively on the corrected time–frequency representation by identifying local maxima that exceeded a noise threshold defined as two times the standard deviation of the residual power, which was recomputed on each iteration. For each detected peak, a two-dimensional Gaussian was fitted around the peak, and its FWHM time and FWHM frequency was estimated to characterise burst duration and bandwidth. For each detected burst, peak frequency, peak time, and FWHM in time and frequency were extracted. The burst waveform was extracted from the preprocessed EEG signal by selecting a 260 ms time window centred on the detected burst peak. The waveform phase was aligned to ensure consistent temporal alignment of the burst trough, after which the segment was mean-centred and polarity-adjusted to ensure a consistent negative deflection before being stored for further analysis.

Burst-feature analyses were conducted at the subject level following within-subject normalisation. For each feature (beta-burst peak amplitude, peak time, peak frequency, FWHM_freq, and FWHM_time), burst-level values were first pooled across motor execution (ME) and motor imagery (MI) conditions and were z-normalised within each participant (z = (x − μ) / σ). A two-way repeated-measures analysis of variance (rmANOVA) with Task (ME, MI) and Force (10%, 25%, 50%, 75%) were performed on normalised burst parameters. F-statistics and associated p-values were computed for the main effects and interaction. Effect sizes were quantified using partial eta squared (η_p_ ^2^). Post hoc pairwise comparisons were conducted separately within each task using two-tailed paired-sample t-tests. Pairwise comparisons were performed between 10% and 25%, 10% and 50%, and 10% and 75% force levels. The resulting p-values were corrected for multiple comparisons using the Benjamini–Hochberg false discovery rate (FDR) procedure. Statistical significance was set at α = 0.05.

#### Beta-burst waveform motif analysis

After detecting all beta bursts from ME tasks (N = 25402 bursts, X ∈ R^25402×130^), to minimise the influence of amplitude outliers while preserving waveform morphology, burst matrices were normalised within each participant using robust scaling (median-centred, interquartile range). Principal component analysis (PCA) was applied to the pooled burst matrix. The number of retained components was determined using a cumulative variance threshold of 99%, ensuring near-complete reconstruction of waveform variability. To identify components most strongly modulated by motor execution, we computed an event-related modulation index (I_m_ score) for each principal component (PC):

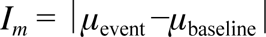

where the Baseline window is −1 to 0 s, and the Event window is 2 to 6 s. The I_m_ score quantified the absolute difference between baseline and event-period PC scores across bursts. PC1 was excluded from selection to avoid temporal skew in the bursts (Papadopoulos, Darmet, et al., 2024; Papadopoulos, Szul, et al., 2024; Szul et al., 2023). The six components with the highest Im scores (excluding PC1) were then selected for motif analysis. For each selected principal component, burst waveforms were separated according to their projection scores. Two kernels were derived per component: a negative kernel (mean waveform of bursts ≤ 5th percentile) and a positive kernel (mean waveform of bursts ≥ 95th percentile). This yielded 12 burst motifs (6 PCs × 2 polarities). Each kernel represents a characteristic waveform pattern underlying task-modulated beta bursts. These kernels were subsequently used as data-driven templates to quantify motif-specific beta dynamics. Preprocessed EEG epochs were band-pass filtered to the beta range (13–30 Hz) using a 4th-order zero-phase Butterworth filter. Each trial was convolved with every kernel maintaining the same data points after convolution. Then we calculated the amplitude envelope then squared it. To obtain a robust reference, the median of the baselines (−0.8 to 0 s) were first computed for each trial across all force conditions and kernels. Baseline correction was performed as [(E(t)−E_baseline_)/E_baseline_ ] × 100. The percentage signal change was then calculated at the single-trial level. Following baseline normalisation, each trial’s convolved time course was temporally aligned with its behavioural reference time, defined as the moment participants reached the target force level. A fixed window spanning −1 to 6 s relative to this alignment point was extracted. Within each force condition, aligned trials were averaged and then smoothed with a Gaussian kernel (σ = 20 samples i.e 40 ms). To quantify force-dependent modulation, a time-resolved linear regression analysis was performed. For each time point, neural activity values were pooled across participants and force conditions (10%, 25%, 50%, 75%) and entered into a linear model, with neural activity as the predictor and force level as the outcome variable. Regression coefficients (β) and associated p-values were computed using ordinary least squares (scipy.stats.linregress) to capture the strength and significance of the relationship between neural activity and force.

To account for multiple comparisons across time points within each kernel, false discovery rate (FDR) correction was applied using the Benjamini–Hochberg procedure (α = 0.05). We performed the same procedure for the MI task (N= 24747 bursts, X ∈ R^24747×130^).

## Results

### Motor execution and force maintenance

For the ME trials, Participants maintained the instructed force levels throughout the force-hold period, with mean deviations from the prescribed target force remaining below 2.5% across all conditions (10%: 0.25%; 25%: 0.23%; 50%: 0.92%; 75%: 1.56%). The time required to reach the target force increased systematically with force magnitude (10%: 1.38 ± 0.43 s; 25%: 1.56 ± 0.46 s; 50%: 1.74 ± 0.48 s; 75%: 1.82 ± 0.49 s), whereas the time point until which the target force was maintained remained relatively stable across force levels (10%: 7.06 ± 0.43 s; 25%: 6.96 ± 0.42 s; 50%: 6.99 ± 0.40 s; 75%: 6.94 ± 0.42 s). A one-way analysis of variance confirmed a strong effect of force level on maintained force (F_3, 1390_ > 100, p < 0.001, η_p_ ^2^ = 0.994), demonstrating sustained control across task conditions. This confirmed that, on average, participants were able to reach the instructed force level and maintain it between 1.8 s to 6.9 s. Therefore, for further analysis, we considered the static period for all participants to be between 2s to 6s.

The EMG from the FCR muscle followed the same pattern, with a strong correlation with the applied force levels (**Supplementary Figure 1**). For the MI trials, the same EMG was used to ascertain whether the participants were performing imagery and not actually applying the force during these trials.

### Mu-beta ERD validates motor execution and imagination

Before analysing the beta bursts, we confirmed classical event-related desynchronization in the mu/beta band (8-30Hz) over the left sensorimotor region, which was used as a spatial filter for each participant (see Methods for details; also see **Supplementary Figure 2**). As seen in many studies on voluntary motor execution, we observed mu-beta desynchronization followed by a mu/beta rebound post-relaxation. Similar, albeit weaker, desynchronization was observed during motor imagery. More interestingly, during the period of force execution and imagination, there were periods of beta event-related synchronisation, which hinted that these could be linked to transient burst events at the single-trial level. **Figure 2C** shows the processing steps for extracting sensorimotor beta bursts. On comparing the differences in the distribution of beta-burst parameters across ME and MI, using a permutation-based Kolmogorov–Smirnov (KS) test (10,000 permutations) on subject-wise normalised beta burst parameters, Significant differences in distributions were observed (**Figure 2D)** for peak amplitude (D = 0.1378, p < 0.001), peak frequency (D = 0.0721, p < 0.001), spectral width (D = 0.0578, p < 0.001), and temporal width (D = 0.0494, p < 0.001), indicating systematic modulation of burst characteristics between conditions. In contrast, no significant difference was found for peak timing (D = 0.0076, p = 0.653), suggesting temporal alignment of bursts remains comparable across ME and MI. These results show that while the temporal occurrence of beta bursts is preserved, their spectral and amplitude properties are significantly altered between execution and imagery conditions, consistent with differential engagement of sensorimotor networks.

### Temporal dynamics of mu and beta ERD - force coupling

Time-resolved regression analysis verifies the dissociation between motor execution (ME) and motor imagery (MI) in the encoding of force within sensorimotor rhythms. During ME, both mu band (8–12 Hz) and beta band (13–30 Hz) ERD exhibited a predominantly negative relationship with force level, indicating stronger desynchronization with increasing force (**Supplementary Figure 3)**. In contrast, during MI, regression coefficients fluctuated around zero in both frequency bands and did not survive multiple comparison correction at any time point, indicating an absence of consistent force-dependent modulation.

### Relationship of beta burst spectral features to motor execution and imagination

Based on our pre-registration , we hypothesised that beta-burst amplitude would decrease as participants applied or imagined higher force levels. A two-way repeated-measures ANOVA with factors Task (ME vs MI) and Force (10%, 25%, 50%, 75%) revealed a significant main effect of Task, with beta-burst amplitudes significantly larger during motor imagery than motor execution (F_1,15_ = 32.867, p < 0.001, η_p_² = 0.687; **Figure 3A**). A significant main effect of Force was also observed (F_3,45_ = 6.498, p = 0.001, η_p_² = 0.302), indicating that burst amplitude varied systematically across force levels. Importantly, the Task × Force interaction was significant (F_3,45_ = 3.146, p = 0.034, η_p_² = 0.173), demonstrating that the force-dependent modulation differed between execution and imagery. Post-hoc paired-sample t-tests with Benjamini–Hochberg FDR correction showed significant decreases in beta-burst amplitude during motor execution between 10% and 25% force (t(15) = 2.214, p = 0.043), 10% and 50% force (t(15) = 3.973, p = 0.002), and 10% and 75% force (t(15) = 4.281, p = 0.002). In contrast, no pairwise differences were observed in motor imagery (all p ≥ 0.366).

**Figure 3.**
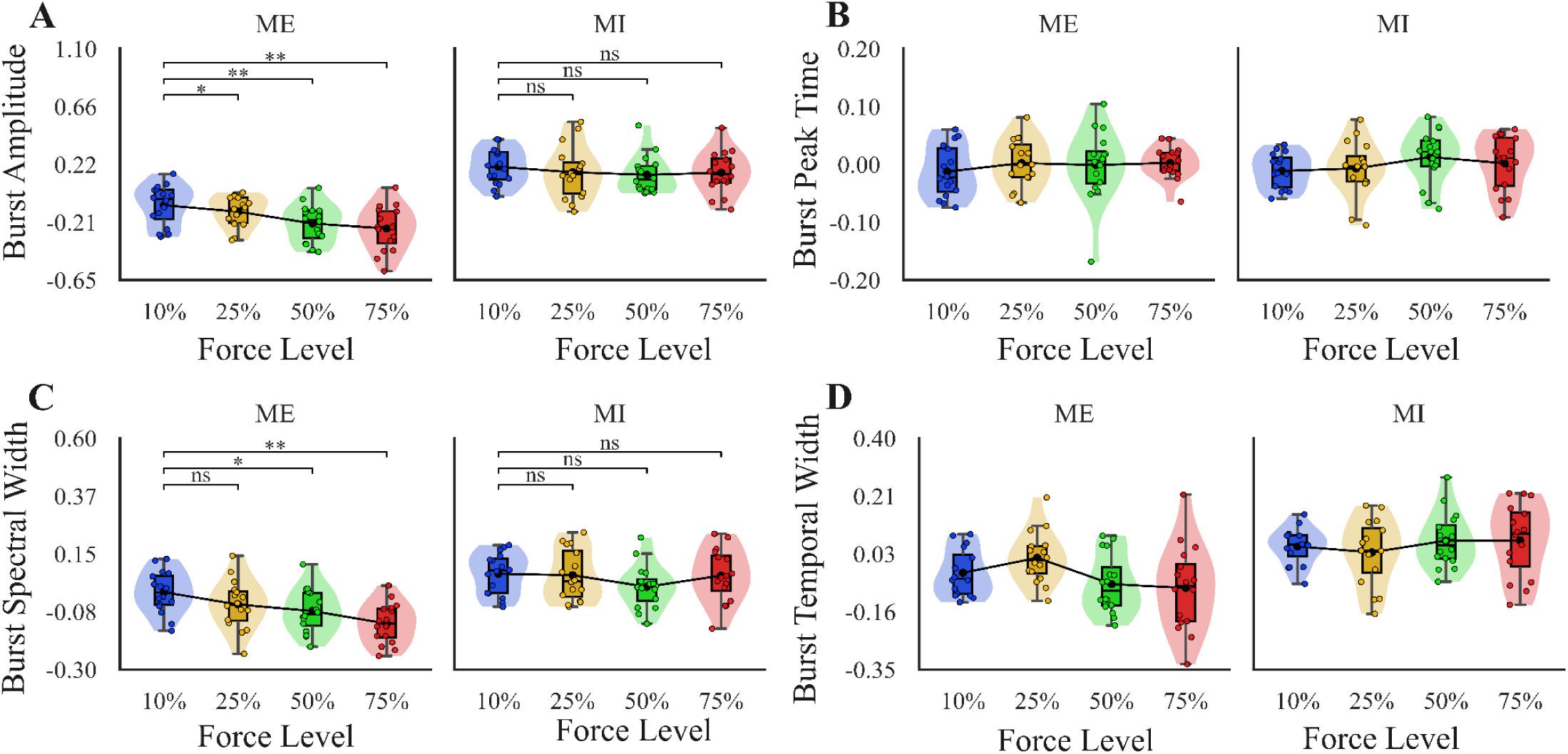
Force-dependent modulation of beta-burst parameters during motor execution (ME) and motor imagery (MI). Violin–box plots show participant-wise distributions across four force levels (10%, 25%, 50%, 75%). Dots represent individual participants, boxplots indicate median and interquartile range, and black lines connect condition means. Pairwise comparisons (10% vs 25%, 50%, 75%) were performed using paired-sample t-tests with Benjamini–Hochberg false discovery rate (FDR) correction; significance is indicated as *p < 0.05, **p < 0.01, ns = not significant. **(A)** Burst amplitude z-scored, **(B)** Burst peak time z-scored, **(C)** Burst spectral width z-scored, **(D)** Burst temporal width z-scored

**Figure 4.**
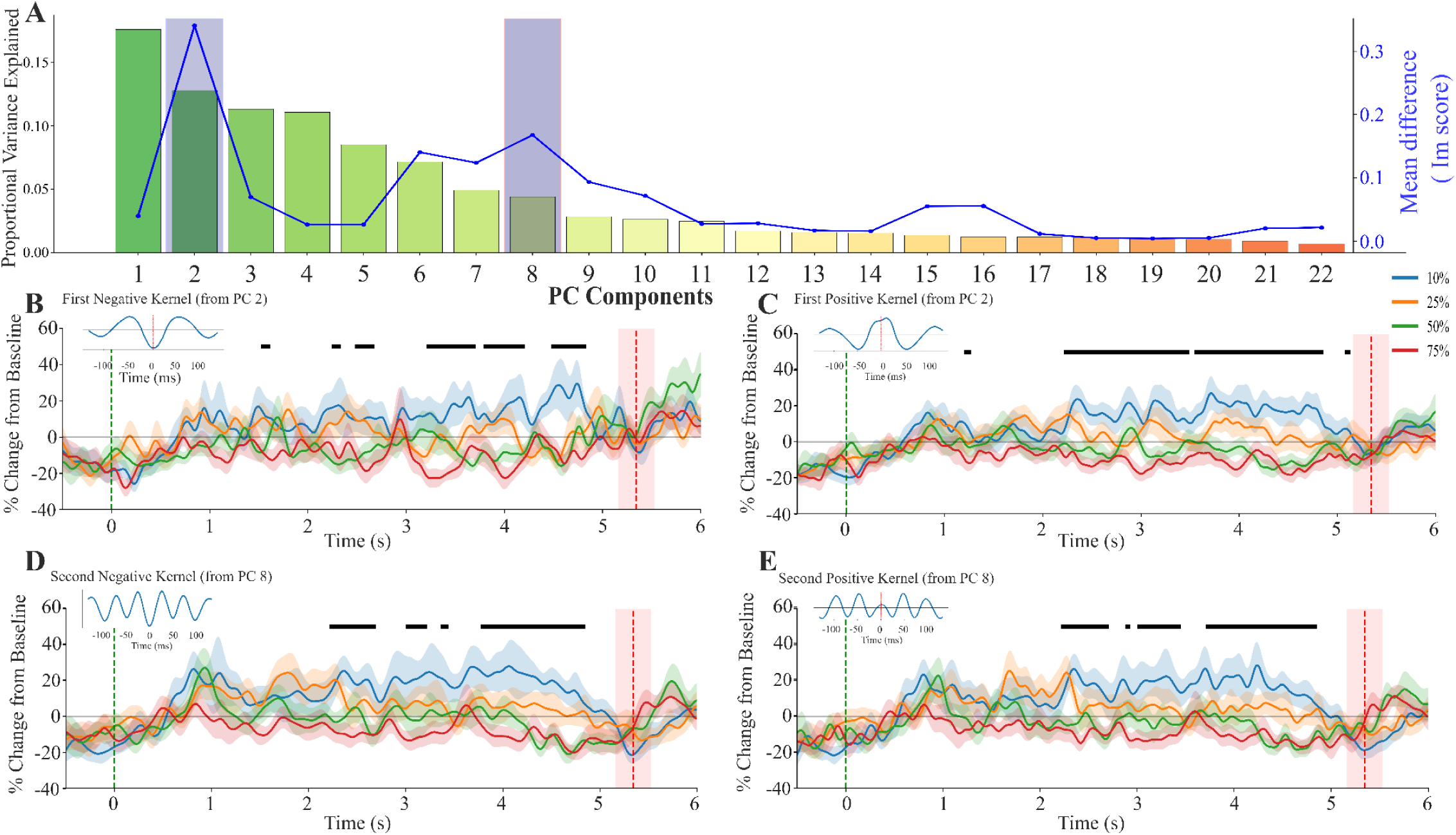
Force-dependent modulation of beta-burst waveform motifs during motor execution. **(A)** Principal component (PC) selection for burst motif extraction. Bars indicate the proportion of variance explained by each PC. The blue line represents the event-related modulation index (Im score), computed as the absolute difference of PC score between baseline (−1 to 0 s) and event (2–6 s) periods. Shaded regions indicate components with the highest modulation values selected for motif analysis (excluding PC1). **(B–E)** Time-resolved motif activation (% change from baseline) for the given kernels (mean ± SEM across participants) across four force levels (10%, 25%, 50%, 75%). Plots are aligned to the reach time (green dashed line at 0 s), defined as the time participants reached the target force level. The red dashed vertical line denotes the mean time till which the participants maintained the force level, with the shaded area indicating ±1 SD. Black horizontal bars denote FDR-corrected time intervals where the regression coefficient (β₁) is significantly less than zero (p < 0.05). Regression coefficient (β₁) variation with time shown in figure 6. For the remaining 8 convolved kernel plots, refer to the **Supplementary** Figure 4.

**Figure 5.**
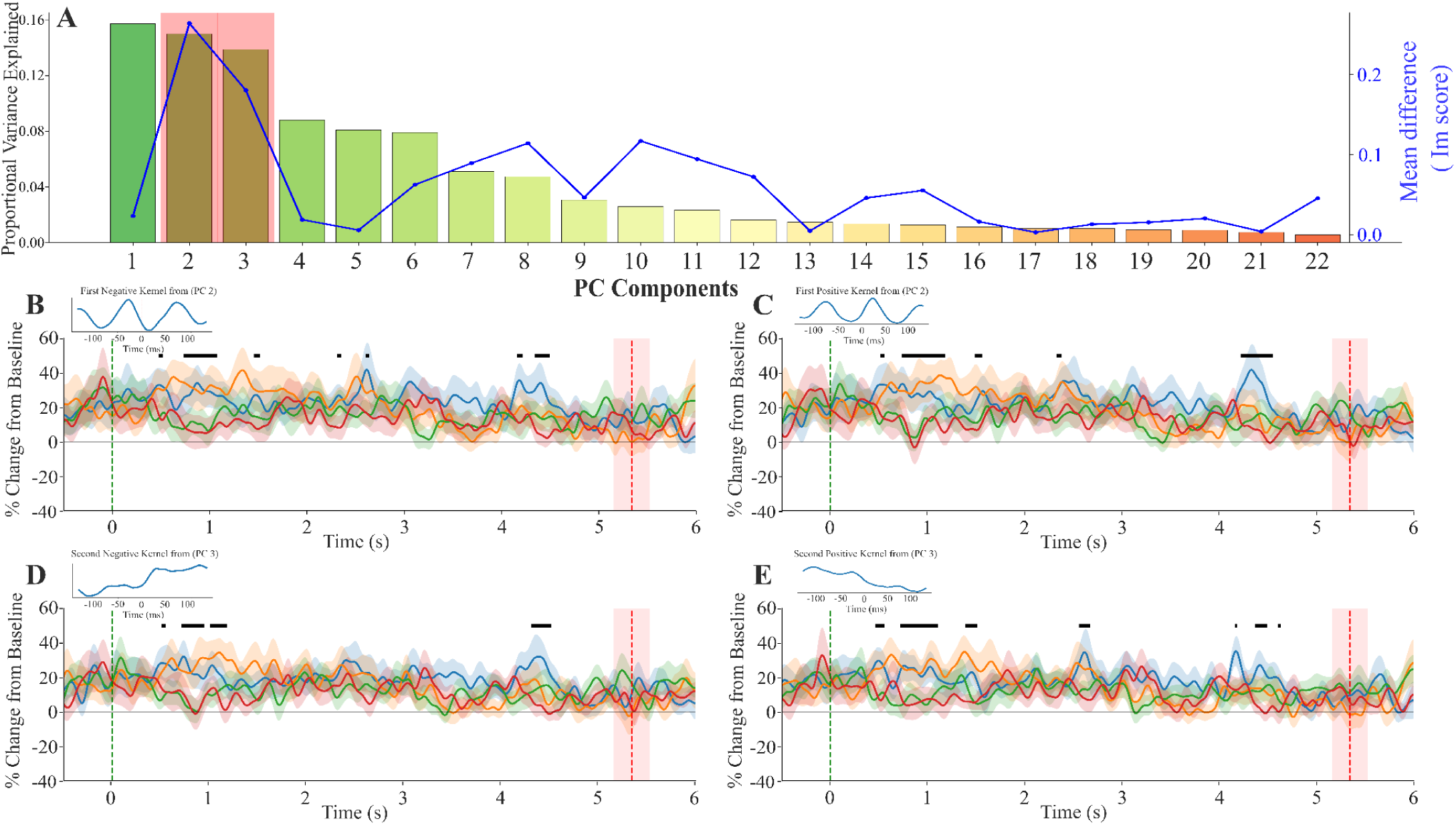
Force-dependent modulation of beta-burst waveform motifs during motor imagery. **(A)** Principal component (PC) selection for burst motif extraction. Bars indicate the proportion of variance explained by each PC. The blue line represents the event-related modulation index (Im score), computed as the absolute difference between baseline (−1 to 0 s) and event (2–6 s) periods. Shaded regions indicate components with the highest modulation values selected for motif analysis (excluding PC1). **(B–E)** Time-resolved motif activation (% change from baseline) for the given kernels (mean ± SEM across participants) across four force levels (10%, 25%, 50%, 75%). Plots are aligned to the reach time (green dashed line at 0 s), defined as the moment participants reached the target force level. The red-dashed vertical line denotes the mean time until which the participants maintained the force level during the motor execution task, with the shaded area indicating ±1 SD. Black horizontal bars denote uncorrected time intervals where the regression coefficient (β₁) is significantly less than zero (p < 0.05)(uncorrected). For the remaining 8 convolved kernel plots, refer to the **Supplementary** Figure 5.

Our next hypothesis was that average beta-burst latency would increase with increasing force levels, reflecting delayed burst occurrence under higher force levels. However, a two-way repeated-measures ANOVA revealed no significant main effect of Force (F_3,45_ = 0.839, p = 0.480, η_p_² = 0.053, **Figure 3B** ), indicating that burst timing did not systematically vary across force conditions. There was also no significant main effect of Task (F_1,15_ = 0.022, p = 0.885, η_p_² = 0.001), suggesting comparable burst latencies between motor execution and motor imagery. The Task × Force interaction was likewise non-significant (F_3,45_ = 0.324, p = 0.808, η_p_² = 0.021), indicating that force-dependent modulation of burst latency did not differ between tasks

In addition to the pre-registered hypothesis, we also did an exploratory analysis on the beta-burst spectral width (FWHM frequency), temporal width (FWHM time), and peak frequency to further characterise task-dependent differences in motor execution (ME) and motor imagery (MI). For spectral width, a two-way repeated-measures ANOVA revealed significant main effects of Task (F_1,15_ = 30.324, p = 0.0001, η_p_² = 0.669, **Figure 3C**) and Force (F_3,45_ = 4.669, p = 0.0063, η_p_² = 0.237) but no significant Task × Force interaction (F_3,45_ = 2.061, p = 0.1188, η_p_² = 0.121), indicating broader bursts during MI and systematic force-related modulation. Post-hoc tests showed that spectral width increased with force during ME (10% vs 50%: p = 0.024; 10% vs 75%: p = 0.003), whereas no pairwise effects were observed during MI. For temporal width, there was a significant main effect of Task (F_1,15_ = 9.893, p = 0.0067, η_p_² = 0.397, **Figure 3D**) and a significant Task × Force interaction (F_3,45_ = 3.830, p = 0.0159, η_p_² = 0.203), but no main effect of Force (F_3,45_ = 0.775, p = 0.514, η_p_² = 0.05); however, post-hoc comparisons did not survive FDR correction within either task. Finally, peak frequency showed no significant main effects of Task or Force, nor an interaction (all p ≥ 0.078). Collectively, these findings indicate robust task-related differences in burst morphology, particularly in burst amplitude and spectral width, and suggest that beta-burst properties are differentially organised between execution and imagery, with implications for feature selection in brain–computer interface applications.

Finally, in accordance with our pre-registeration, we also assessed whether beta-burst parameters improve force-level prediction using hierarchical general linear models and complementary Bayesian model comparison. In the motor execution task, the baseline model (mu and beta power) significantly predicted force (F₂,₆₁ = 9.73, p = 0.0002, R² = 0.242, AIC = 580.62), whereas inclusion of beta-burst amplitude and timing produced only a marginal increase in explained variance (R² = 0.291, AIC = 580.29) without significant improvement (LR χ²(2) = 4.33, p = 0.115; ΔAIC = −0.33). Bayesian analysis similarly favoured the baseline model (BF_10_ = 1.00, R^2^ =0.242), while addition of beta-burst amplitude provided modest improvement in model evidence (BF_10_ =1.27, R^2^ = 0.282). Inclusion of beta-burst timing did not improve model performance, as models containing burst timing showed lower evidence relative to the baseline (BF_10_ =0.38, R^2^ =0.248). In the motor imagery task, neither the baseline (F₂,₆₁ = 0.27, p = 0.763, R² = 0.009, AIC = 597.78) nor the full model (F₄,₅₉ = 1.52, p = 0.209, R² = 0.093, AIC = 596.08) was significant, and model improvement was not supported (LR χ²(2) = 5.70, p = 0.058; ΔAIC = −1.70).

These findings suggest that conventional regression frameworks based on trial-averaged features may not fully capture the functional contribution of beta-burst dynamics to force modulation. In particular, summarising bursts through averaged amplitude or timing metrics likely obscures their transient and temporally structured nature (Szul et al., 2023). To address this limitation, we therefore adopt a time-resolved, motif-based waveform analysis that preserves the temporal organization of beta activity, providing a more sensitive approach for characterizing force-dependent neural dynamics.

### Beta Burst Waveform Analysis

During motor execution (ME), robust force-dependent modulation of beta-burst waveform motifs was observed. Significant effects (FDR-corrected, p < 0.05) were primarily concentrated in the stable-force period. Force-related differences emerged and persisted over time. Higher force contrasts (10% vs 50% and 10% vs 75%) produced the strongest separations, indicating a robust and graded relationship between beta-burst features and force level during active movement.

In contrast, during motor imagery (MI), beta-burst waveform motifs showed weak and inconsistent modulation with force level. Significant effects (uncorrected, p < 0.05) were sparse, short-lived, and distributed across time, without clear clustering in the stable imagery period. There was no sustained force-dependent separation, and trajectories across force levels largely overlapped.

### Beta-burst motifs exhibit stronger relation to force than classical power measures

We can observe from the both time-resolved regression analysis of the ME task that beta burst derived features exhibit a stronger and more consistent relationship with force levels compared to mu-ERD (**Figure 6**). Across the movement epoch, beta burst regression coefficients (β₁) showed larger magnitude deviations, indicating enhanced sensitivity to graded force modulation. In contrast, mu ERD demonstrated relatively weaker associations with force, with smaller regression slopes.

**Figure 6:**
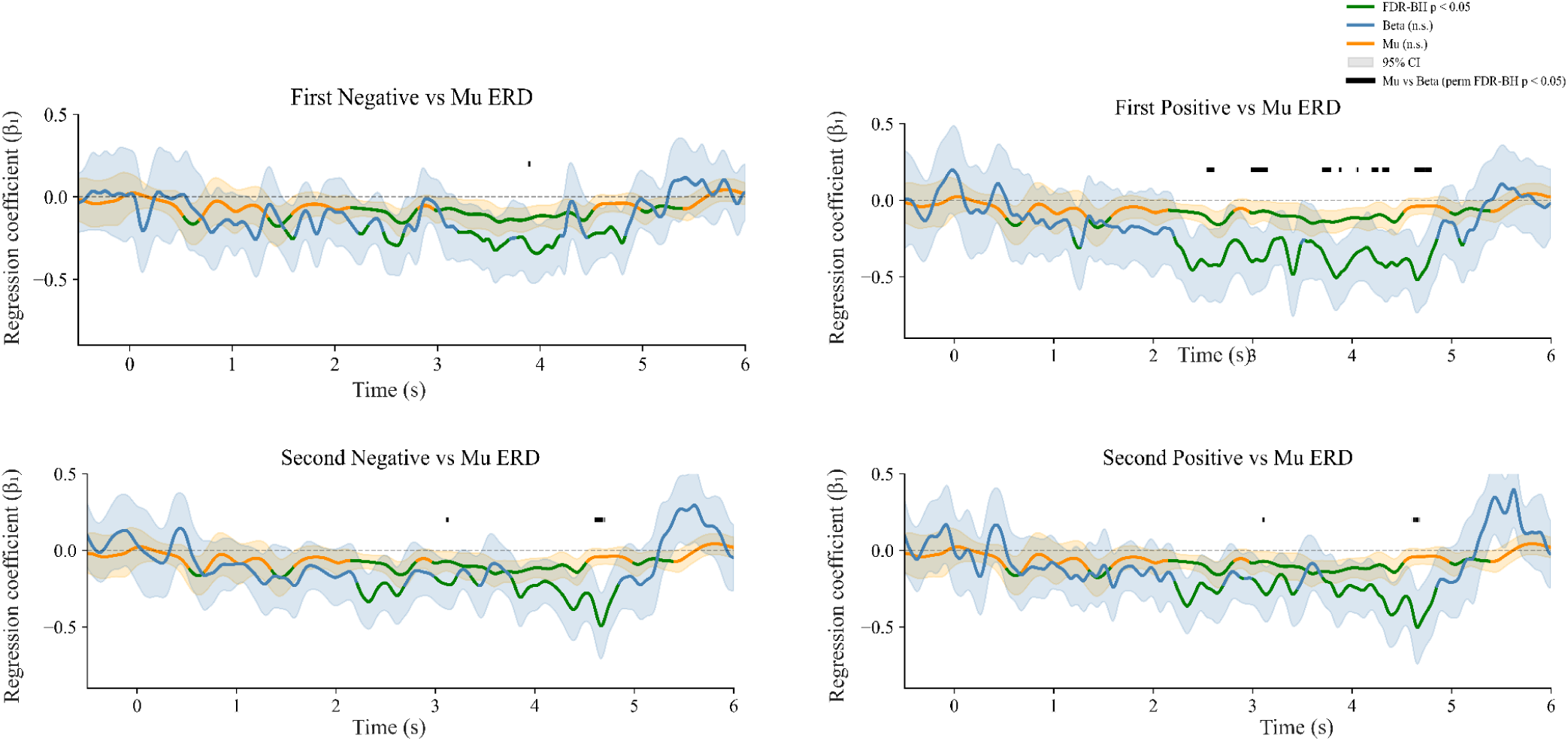
Time-resolved regression coefficients (β₁; mean ± 95% confidence intervals) relating neural activity to force magnitude for different beta-burst motifs and mu ERD during motor execution. Green traces indicate time points where the regression coefficient (β₁) is significantly (FDR corrected ) less from zero (p < 0.05), reflecting significant encoding of force magnitude, whereas blue (beta motifs) and orange (mu ERD) traces denote non-significant intervals. Black horizontal bars denote time intervals where beta-burst–derived regression coefficient significantly less than mu ERD regression coefficient (p < 0.05). Signals are aligned to reach time (0 s).

Permutation-based statistical comparisons further confirmed these observations. Significant differences between beta burst and mu ERD representations were observed over extended time intervals, as indicated by the horizontal significance bars. These results demonstrate that beta burst features capture force-related neural dynamics more robustly than traditional mu power measures, particularly during sustained motor output phases.

Importantly, while both signals exhibited periods of significant force encoding, beta bursts provided a more temporally stable and higher-amplitude representation, suggesting that transient burst events may serve as a more precise neural marker of motor control than continuous oscillatory power measures such as mu ERD.

In contrast to the movement execution (ME) findings, the motor imagery (MI) condition revealed a comparatively weaker dissociation between beta burst features and mu power representations. Although beta burst–based time-resolved regression exhibited relatively higher β₁ values than mu ERD across portions of the imagery epoch, these differences did not reach statistical significance under permutation-based inference.

## Discussion

Force production is generally accompanied by the classic mu-beta ERD over sensorimotor cortex, with stronger forces producing higher and more sustained desynchronisation (Alegre et al., 2004; Pfurtscheller et al., 1996; Torrecillos et al., 2018). In accordance with the established results we saw a similar trend was observed during MI, but it was much weaker and less reliable than during ME (Duann & Chiou, 2016; Lotze & Halsband, 2006; Munzert et al., 2009). To establish alternative method and move beyond the classic trial averaged frequency band power based classification, we explored the role of beta bursts in decoding different force levels (Echeverria-Altuna et al., 2022; Papadopoulos, Szul, et al., 2024) using beta burst parameters and burst-waveform motifs. At the burst level, we found a clear dissociation between ME and MI burst features (Van der Lubbe et al., 2021). Beta bursts during motor execution scaled systematically with force, showing reduced amplitude and distinct waveform modulation as force increased, whereas imagery lacked reliable force-dependent effects. Notably, burst timing relative to stimulus onset was the same across tasks, suggesting preserved temporal organisation, but burst amplitude and spectral features differentiated execution from imagery, pointing to partially overlapping yet functionally distinct mechanisms. This means that we need different, tailored approaches to decode motor intent from imagination, rather than merely treating MI decoding as an extension of ME decoding (Duann & Chiou, 2016; Lotze & Halsband, 2006; Munzert et al., 2009). Modulation of motif-convolved EEG data during the stable-force window further supports a tight coupling between beta-burst dynamics and actual motor output. We observed a similar modulation in MI, although it did not cross the statistical threshold. The weaker imagery effects may reflect both neurophysiological differences and methodological factors, including limited MI training and the absence of real-time feedback, which could have reduced the precision of imagined force scaling (Behrendt et al., 2021; Hwang et al., 2009; Lebon et al., 2010). Currently, efforts to decode continuous force from EEG data primarily operate at the macroscopic level, focusing on the sustained, trial-averaged power modulation of specific frequency bands such as mu (8–12 Hz), beta (13–30 Hz), and low-gamma (30–45 Hz) as well as movement-related cortical potentials (Chakarov et al., 2009; Fry et al., 2016; Haddix et al., 2021; Kristeva-Feige et al., 2002; Peter et al., 2022; Torrecillos et al., 2018).

To get continuous force decoding, researchers typically extract continuous band-power amplitudes or spatial features and feed them into regression models or classifiers, operating under the conventional assumption that sensorimotor rhythms are smoothly sustained, amplitude-modulated oscillations (Haddix et al., 2021). However, our findings demonstrate that, in addition to continuous mu and beta power, sensorimotor beta bursts can be used to decode different force levels. The transient nature of these oscillations makes them an interesting signature to decode force at single-trial levels. The shape of the burst waveform that separates the different force levels during execution and imagery could lead to individual-specific neural decoding strategies. Burst waveforms can functionally denote distinct motifs that serve as proxies for the transient activation of specific underlying neural circuits, meaning that different burst shapes signify distinct computational operations (Little et al., 2019; Lundqvist et al., 2024; Rayson et al., 2026; Szul et al., 2023). In the context of motor control, these motifs are highly behaviorally relevant because they are selectively modulated according to given task demands with certain motifs decreasing in rate to drive pre-movement desynchronisation and others increasing to drive post-movement synchronisation, thereby dynamically adapting continuous motor performance, peak movement velocity, and reaction times (O’Keeffe et al., 2020; Rayson et al., 2026; Szul et al., 2023; Torrecillos et al., 2018). At the cellular level, this varied bursting activity is intimately tied to neuronal spiking; cortical beta bursts are associated with the active suppression of informative neuronal spiking and gamma activity (Lundqvist et al., 2018). Existing BCI systems for force decoding have mostly relied on trial-averaged ERD/S of sensorimotor mu and beta rhythms to infer discrete motor states or coarse grip categories (Edelman et al., 2016; Pfurtscheller & Lopes da Silva, 1999). Although viable for categorical classification, averaged band power evolves slowly, saturates across intermediate force levels, and collapses trial-by-trial variability into a smooth envelope that cannot track the rapid modulations required for proportional control (Fry et al., 2016; Little et al., 2019) leaving BCIs ability to gate movement initiation but unable to resolve graded force intent. Our results show that sensorimotor beta bursts offer a way to bridge this gap. Rather than treating the EEG as a sustained oscillation whose mean power encodes state, isolating discrete burst events reveals a temporally precise, trial-varying neural signature whose rate, timing, and waveform morphology are tightly coupled to motor output (Echeverria-Altuna et al., 2022; Little et al., 2019; Rayson et al., 2026; Shin et al., 2017; Szul et al., 2023; Torrecillos et al., 2018). Our results in accordance with the recent work demonstrates that using specific burst waveform motifs as data-driven temporal filters yields decoding accuracies and Information Transfer Rates that surpass standard band-power models (Papadopoulos, Szul, et al., 2024; Szul et al., 2023), and this could also unlock sensitivity to graded force levels that averaged power simply cannot resolve. Adopting this burst-centric framework thus represents not an incremental refinement but a necessary paradigm shift for achieving continuous, proportional force decoding in next-generation non-invasive neuroprosthetics.

Achieving highly dexterous control of neuroprosthetics requires the precise decoding of individual finger movements and continuous force outputs. Invasive and semi-invasive methods, such as electrocorticography (ECoG), have demonstrated exceptional capabilities in this domain due to their high spatiotemporal resolution; researchers have successfully utilised low-frequency local motor potentials (LMPs) and high-gamma band activity to accurately predict 3D fingertip trajectories and classify individual finger flexions (Acharya et al., 2010; Nakanishi et al., 2014; Volkova et al., 2019). Building on these spectral advantages, ECoG-based systems have further demonstrated online, real-time individual prosthetic finger control using native high-gamma cortical representations without arbitrary remapping, and have decoded up to three distinct hand gestures at latencies below 500 ms using recurrent neural networks, establishing ECoG as the closest non-spiking bridge between single-unit recordings and scalp-level signals (Hotson et al., 2016; Pan et al., 2018). Noninvasive scalp electroencephalography (EEG) has historically struggled to differentiate highly overlapping sensorimotor tasks, such as representations of individual digits or different force levels, due to signal attenuation and volume conduction. The coarse spatial resolution of scalp electrodes means each sensor integrates activity from thousands of spatially overlapping neuron populations. EEG signals exhibit fundamentally degraded discriminability, yielding accuracies of 43–60% for individual-finger classification tasks, whereas ECoG achieves over 91% on identical paradigms (Lee et al., 2022; Liao et al., 2014). Our results provide a way to overcome this by adopting burst-centric framework wherein transient, discrete beta burst events replace sustained band-averaged power as the primary decoding feature, not only promise to enhance the accuracy and speed of non-invasive neuroprosthetics but also open entirely new avenues for decoding cortical intent from peripheral neural interfaces. Unlike classical ERD/S measures that smooth over trial-to-trial variability, the timing, waveform morphology, and rate of sensorimotor beta bursts encode task-specific information at the single-trial level, offering a richer representational substrate for graded and continuous motor intent decoding (Papadopoulos et al., 2022; Papadopoulos, Szul, et al., 2024).

In conclusion, relying solely on sustained mu and beta-band power is insufficient for accurate, continuous force decoding. Our results show that sensorimotor beta bursts, characterised by diverse and functionally distinct waveform motifs, offer a richer neural signature that tightly couples to actual motor output. Furthermore, the clear dissociation in burst dynamics between motor execution and motor imagery indicates that MI is not just a weaker version of physical movement, necessitating the development of distinct, motif-based decoding modalities for mental simulation. Embracing this burst-centric paradigm provides a critical pathway to engineering the next generation of highly dexterous, force-sensitive brain-computer interfaces.

## Author Contributions

Md Shaheen Perwez: Conceptualization, Data curation, Formal analysis, Investigation, Methodology, Validation, Visualization, Writing – original draft, Writing – review & editing.

James J. Bonaiuto : Supervision, Validation, Writing – review & editing Bhivraj Suthar : Conceptualization, Project administration, Supervision. Vignesh Muralidharan : Conceptualization, Funding acquisition, Investigation, Methodology, Project administration, Resources, Supervision, Validation , Writing – original draft, Writing – review & editing

## Acknowledgments

The author used AI assistance tools for limited support in code development and language editing of the manuscript. These tools did not contribute to the study design, data analysis, interpretation of results, or scientific conclusions. All AI assisted outputs were critically reviewed and edited by the author, who retains full responsibility for the content.

