## Supplementary Figures for "Investigating sensorimotor beta burst dynamics as a robust biomarker for graded force modulation in humans"

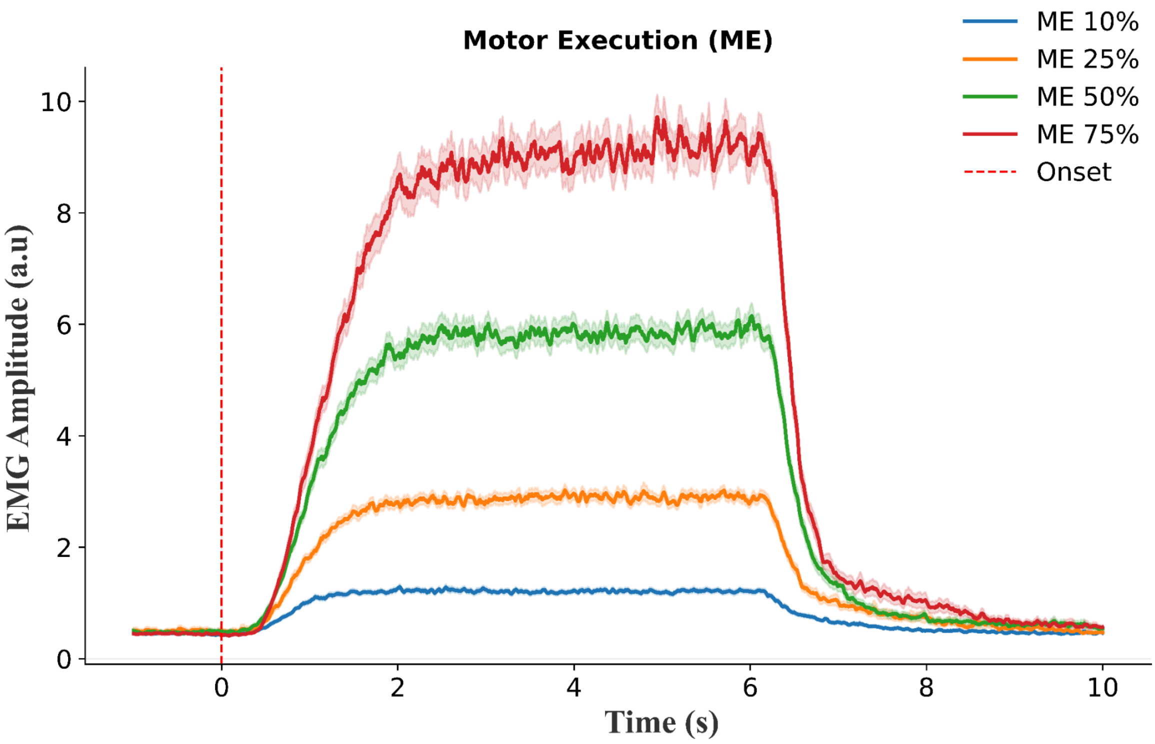


**Supplementary Figure 1. Force-dependent modulation of EMG amplitude during sustained isometric contraction.**
Grand-averaged rectified EMG rms amplitude time courses for four force levels (10%, 25%, 50%, and 75% of maximum voluntary contraction) are shown time-locked to event onset (0 s; red dashed line). Signals were band-pass filtered and smoothed using a moving-average window prior to averaging across trials. Shaded regions represent ±1 SEM across epochs. Following event onset, EMG amplitude increased in a graded, force-dependent manner, with higher contraction levels exhibiting larger and more sustained envelope amplitudes.

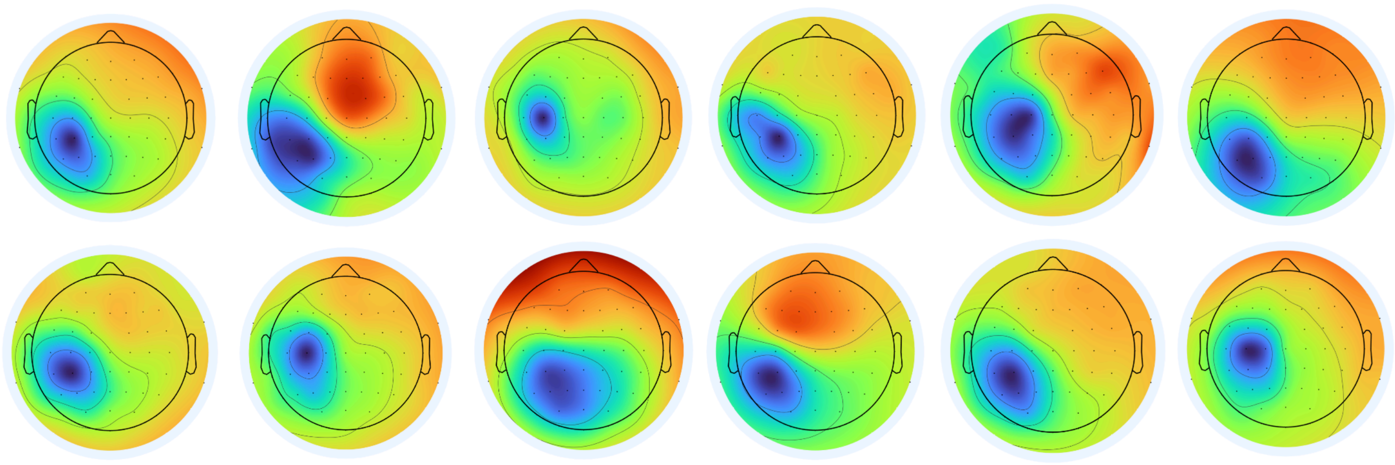

**Supplementary Figure 2:**Spatial topographies of independent components (ICs) selected for analysis from 12 participants (out of 16 participants included in the study).

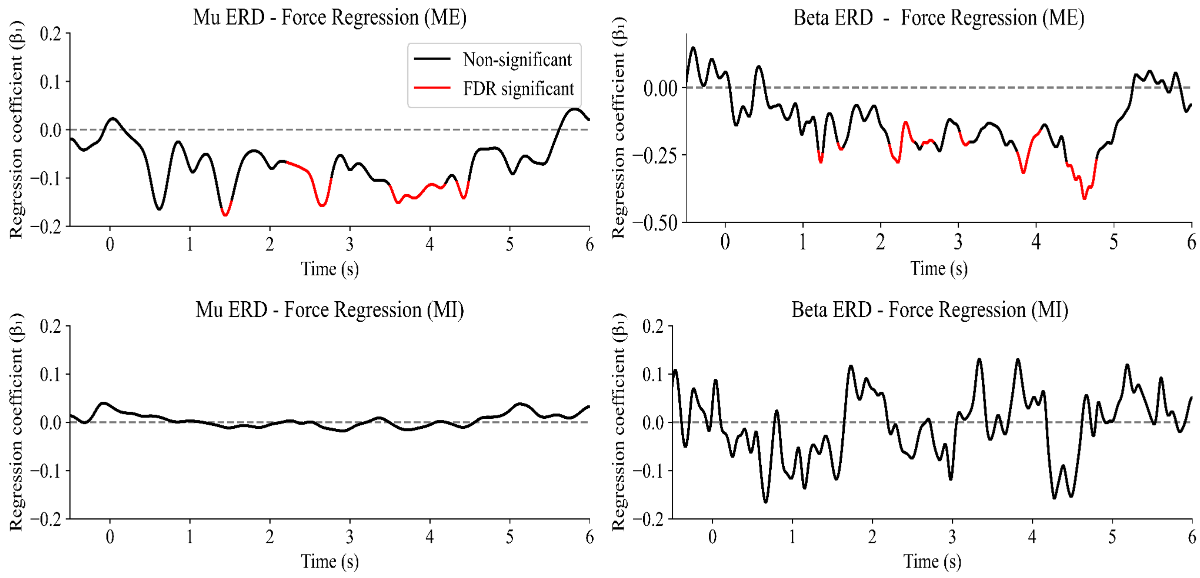


**Supplementary Figure 3:** Time-resolved linear regression between mu and beta band ERD and force level for motor execution (ME) and motor imagery (MI). The regression coefficient ($\beta_1$) is shown as a function of time for mu band (8–12 Hz; left column) and beta band (13–30 Hz). Red segments indicate time points that survived false discovery rate (FDR) correction ($p < 0.05$), while black segments denote non-significant intervals. A sustained negative relationship between ERD and force is observed during ME in both mu and beta bands. In contrast, MI shows weak, non-significant modulation over time in both frequency bands, suggesting limited force-related encoding in the absence of overt movement.

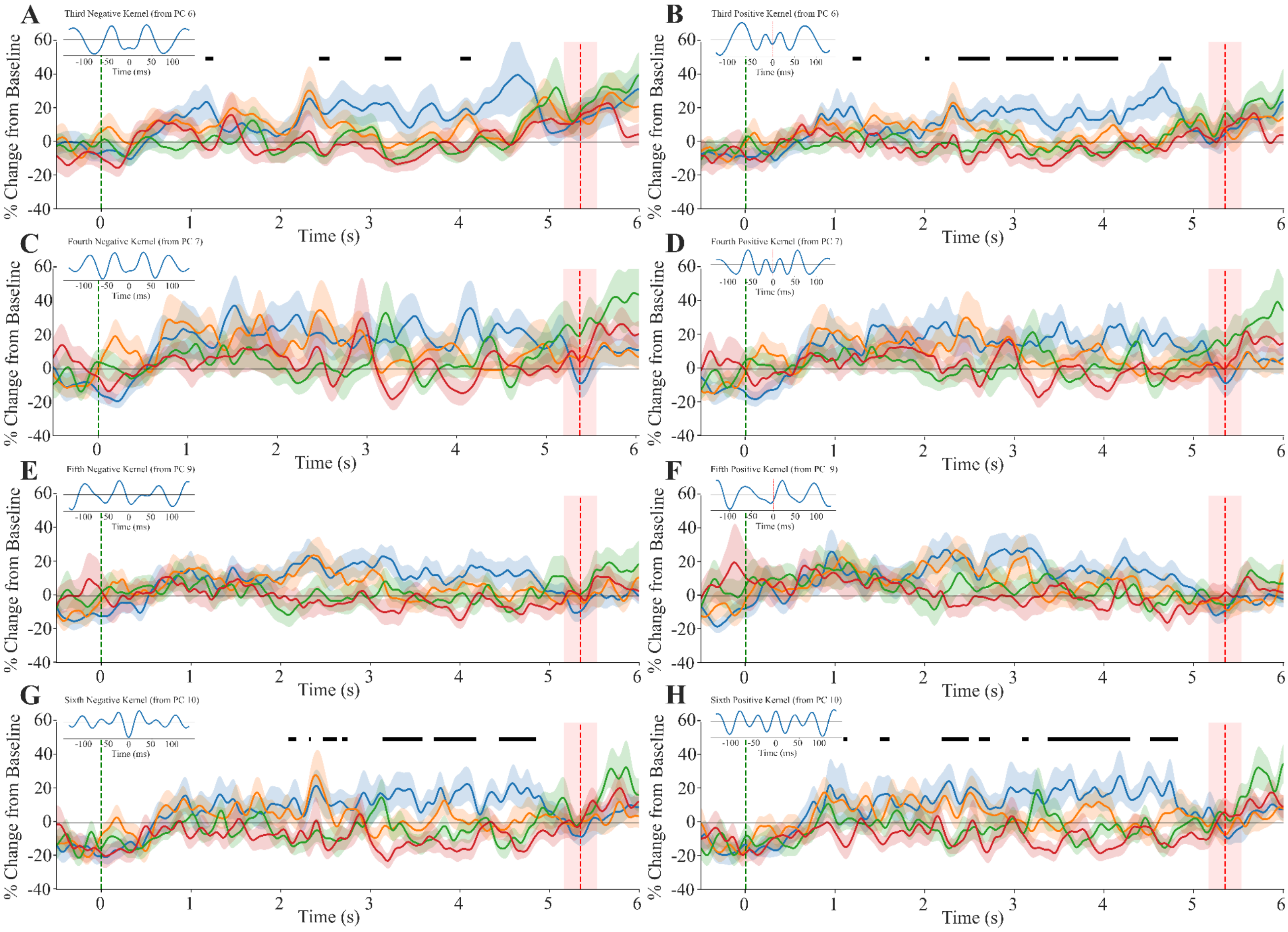

**Supplementary Figure 4:** Force-dependent modulation of beta-burst waveform motifs in ME.
Time-resolved motif activation (% change from baseline; mean ± SEM across participants) for all extracted burst kernels (first to sixth, negative and positive phases) across four force levels (10%, 25%, 50%, 75%). Signals are aligned to reach time (green dashed line at 0 s), defined as the moment participants reached the target force level. The red-dashed vertical line indicates the mean duration of force maintenance, and the shaded region represents ±1 SD across participants.
Black horizontal bars denote FDR-corrected statistically significant ($p < 0.05$) time intervals where the regression coefficient ($\beta_1$) is less than zero. Subplot (A) derived using the third negative kernel, Subplot (B) derived using the third positive kernel, Subplot (C) derived using the fourth negative kernel, Subplot (D) derived using the fourth positive kernel, Subplot (E) derived using the fifth negative kernel, Subplot (F) derived using the fifth positive kernel, Subplot (G) derived using the sixth negative kernel and Subplot (H) derived using the sixth positive kernel.

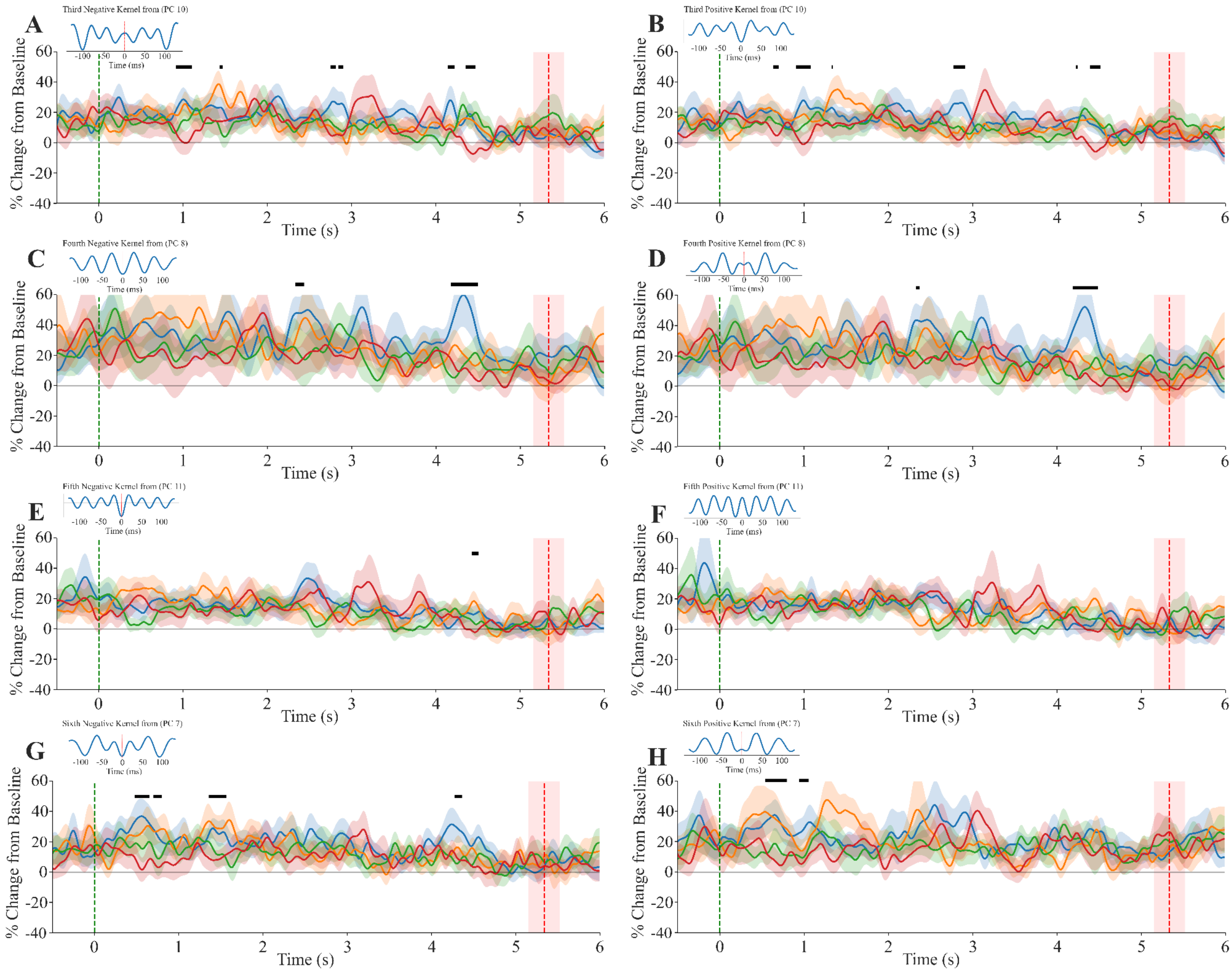


**Supplementary Figure 5:** Force-dependent modulation of beta-burst waveform motifs in MI.
Time-resolved motif activation (% change from baseline; mean ± SEM across participants) for all extracted burst kernels (first to sixth, negative and positive phases) across four force levels (10%, 25%, 50%, 75%). Signals are aligned to reach time (green dashed line at 0 s) as per ME trials, defined as the moment participants reached the target force level. The red-dashed vertical line indicates the mean duration of force maintenance, and the shaded region represents ±1 SD across participants.
Black horizontal bars denote uncorrected statistically significant ($p < 0.05$) time intervals where the regression coefficient ($\beta_1$) is less than zero.
Subplot (A) derived using the third negative kernel, Subplot (B) derived using the third positive kernel, Subplot (C) derived using the fourth negative kernel, Subplot (D) derived using the fourth positive kernel, Subplot (E) derived using the fifth negative kernel, Subplot (F) derived using the fifth positive kernel, Subplot (G) derived using the sixth negative kernel and Subplot (H) derived using the sixth positive kernel.
